# Model reconstruction from Small angle X-ray Scattering data using deep learning methods

**DOI:** 10.1101/691832

**Authors:** Hao He, Can Liu, Haiguang Liu

**Author notes:** **Funding information** National Natural Science Foundation of China (grant No. 11575021, U1530401, U1430237 to Haiguang Liu).

## Abstract

We present an algorithm based on a deep learning method for model reconstruction from small angle X-ray scattering (SAXS) data. An auto-encoder for protein 3D models was trained to compress 3D shape information into vectors of a 200-dimensional latent space, and the vectors are optimized using genetic algorithms to build 3D models that are consistent with the scattering data. The algorithm was implemented using Python with the TensorFlow framework and tested with experimental data, demonstrating capacity and robustness of accurate model reconstruction even without using prior model size information.

**Synopsis:** A deep learning method based on the auto-encoder framework for model reconstruction from small angle scattering data

## 1. Introduction

Small angle X-ray scattering (SAXS) from protein molecules in solution is a powerful technique, providing information on molecular structures and dynamics (Putnam *et al*., 2007; Grant *et al*., 2011; Svergun & Koch, 2003). Because the solution scattering method does not require special treatment for protein molecules, such as crystallization in diffraction measurement or isotope labelling in nuclear magnetic resonance, SAXS experiments can be performed in high-throughput manners (Hura *et al*., 2009). Another major advantage of SAXS experiments is the ability to probe the structure and dynamics in solution, especially when combined with pumping methods to promote conformational changes (Neutze & Moffat, 2012; Kim *et al*., 2012). Time-resolved studies will reveal important information on molecular mechanism for protein functions.

Despite the success in extracting structural information from SAXS profiles, reconstructing high-quality 3D models remains challenging. Several approaches have been proposed and implemented to build 3D density maps from SAXS data, including shape envelope approximation using spherical harmonics functions, polymer chain folding, dummy atom assembly, iterative phasing, and database searching methods. The spherical harmonics function approximation method is fast but limited by resolution (Stuhrmann, 1970; Svergun & Stuhrmann, 1991; Svergun *et al*., 1996). In the Gasbor program, polymers composed of connected beads were used to represent protein molecules, and packing of these polymers was optimized to build 3D models (Svergun *et al*., 2001). Dummy atoms arranged in a 3D lattice were also used for model reconstruction, as implemented in DAMMIN/DAMMIF (Svergun, 1999; Franke & Svergun, 2009). An iterative phase retrieval method was expanded to analyse SAXS data and demonstrate its potentials (Grant, 2018). A database of shapes abstracted from actual protein complexes and efficiently represented using 3D Zernike polynomials was used to quickly retrieve 3D models that match experimental SAXS profiles, as implemented in sastbx.shapeup (Liu, Hexemer *et al*., 2012). A real space representation of a 3D model requires many parameters, such as the position of each bead, which can be described using its coordinates (then 3N parameters are required for a model with N beads). However, the number of parameters required is much greater than the number of free parameters encoded in 1D SAXS profiles. Therefore, prior knowledge must be applied to provide additional constraints for converged reconstructions. For example, the molecular size and the connectivity of the beads are very critical for DAMMIN/DAMMIF. In the case of sastbx.shapeup, the molecular size is de-coupled from the abstracted shapes (Liu, Morris *et al*., 2012), allowing an optimization of the size as a separate parameter. However, the diversity of models is limited by the database. Model reconstruction will be advanced if the following criteria are met: (1) diverse shapes of 3D models can be efficiently represented to cover a broader range than those in structure databases; and (2) SAXS profiles can be computed for each model that can be scaled to arbitrary sizes. We provide a solution to achieve this using an auto-encoder method combined with 3D Zernike representations (Canterakis, 1999; Liu, Morris *et al*., 2012).

Inspired by deep learning methods, real space 3D models were encoded using an auto-encoder neural network to a compressed representation of 200 latent parameters. The protein complexes in the PISA database (Krissinel & Henrick, 2007) were used to generate the training datasets for the auto-encoder. Each complex structure was scaled to fit within a unit sphere then voxelized to a 3D grid of 31×31×31. Because SAXS data often provide low-resolution information that warrants a uniform density approximation for 3D models, we binarized the voxelized objects before auto-encoder training (due to this uniform density approximation, the SAXS data comparison were limited up to q=0.2 Å−1). The testing results show that the shapes represented using 313 voxels with binary numbers can be encoded using a vector of 200 dimensions. The reduction of parameter space allows applying optimization algorithms to improve the model-data fitting. SAXS profiles for 3D voxelized objects were computed using the Zernike expansion method, taking advantage of fast evaluation of theoretical profiles at an arbitrary model radius. If desired, this radius will be coded using an additional parameter and subject to the optimization along with the other 200 parameters. The testing results using experimental data from the SASBDB (Valentini *et al*., 2015) and BIOISIS (Rambo & Tainer, 2011) show that the proposed method can successfully generate 3D models based on SAXS data. The algorithm is implemented to the software, decodeSAXS, whose source code and an associated webserver are available at http://liulab.csrc.ac.cn/decodeSAXS.

## 2. Methods

### 2.1 Training and Testing Datasets

The model dataset is compiled from the PISA structure database, including 60,000 randomly selected models. Each model was first scaled and shifted to fit in a sphere centred at the coordinate origin with a radius of 50 Å. Then, the atomic positions were mapped to a grid of 31 × 31 × 31 in the process of scaling and voxelization. As a result, each model was converted to a voxel object described using a 3D matrix with binary values (see Figure 1a). The 31 × 31 × 31 binary matrix was padded with zeros to a matrix of 32 × 32 × 32 before neural network training.

**Figure 1.**
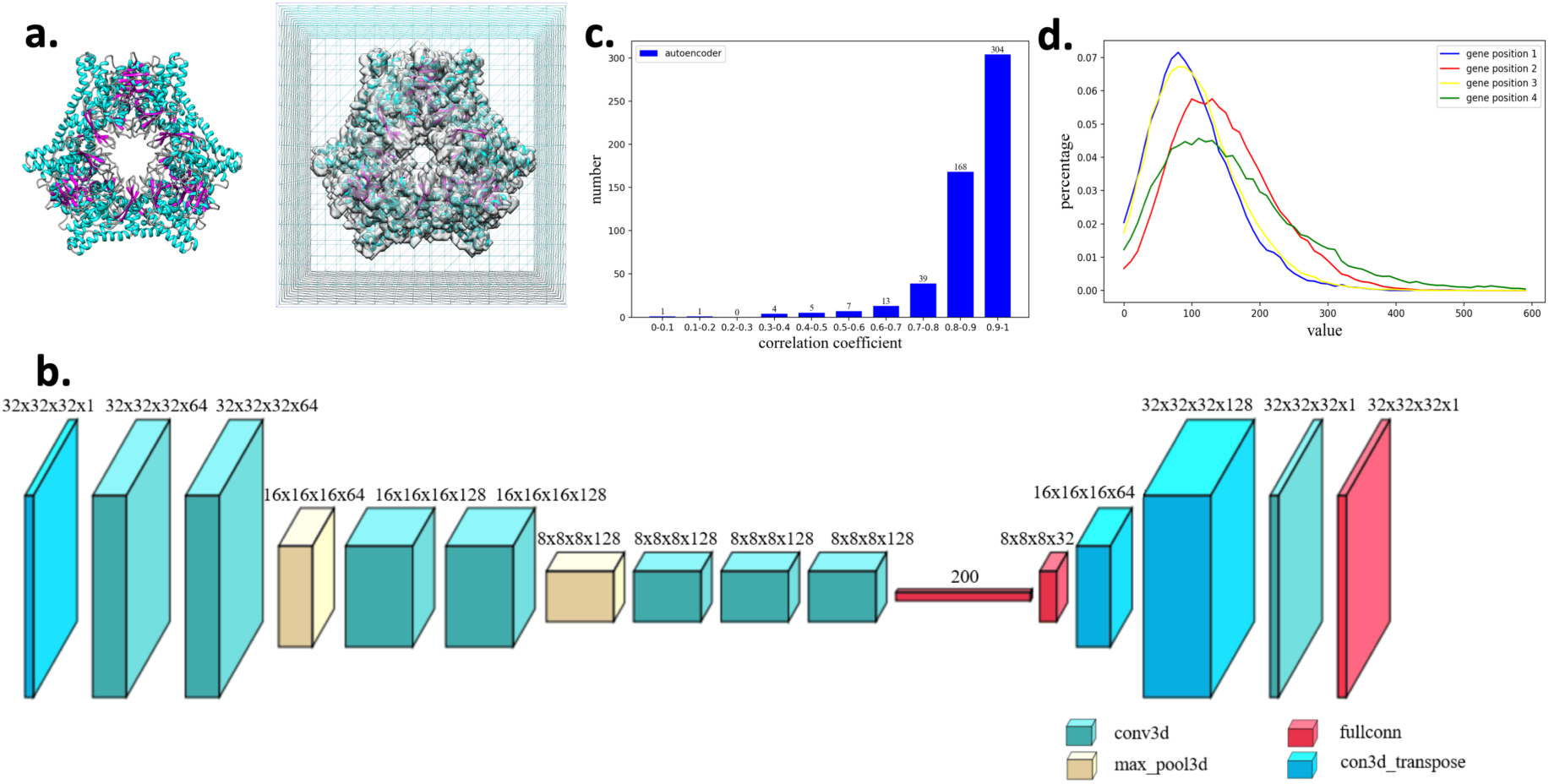
Framework of auto-encoder and its capability in representing 3D models. (**a**) The voxelization of a molecular structure. Left: an atomic model represented in the cartoon representation. Right: the model is mapped on a 3D matrix whose values are binarized depending whether the grids are in the vicinity of any atom of the protein complex. (**b**) The auto-encoder architecture used in this study. The layers and structures are shown in the figure; details can be found in the Methods section. (**c**) The model quality encoded using the trained network and measured using the Pearson correlation coefficients between the models before and after going through the auto-encoder. (**d**) The distribution of encoding parameter values for first four gene positions.

### 2.2 Auto-encoder Neural network architecture

The architecture of the auto-encoder is designed based on the VGG network (Simonyan & Zisserman, 2014). The encoding part of the auto-encoder is composed of 7 convolution layers and 2 pooling layers followed by a dense (fully connected) layer as indicated in Figure 1b. Network training was performed in two stages. During the first stage, the dense layer contains 3,000 variables, and this number is reduced to 200 during the second stage. With this design, the final output is a 3,000-dimensional vector after the first training stage. Among these 3,000 parameters, a significant portion (approximately 90%) of parameters is found to be zeros persistently, indicating that the parameter space for encoding can be reduced. With this observation, the fully connected layer was optimized again with a reduced dense layer of 200 parameters. During the second stage of training, the parameters for convolutional and pooling layers were inherited from the first stage and remained unchanged, except that the parameters for the fully connected dense layer were subjected to optimization. This two-stage training was adapted to ensure fast convergence of the training (we found that if the dense layer was set to a 200-dimension vector during the first stage of training, the loss function does not converge).

The decoding part is relatively simple. The 200-dimension vector is converted to a 4D tensor of size 8 × 8 × 8 × 32, then followed by two deconvolution layers, and finished with a convolution layer to obtain a 32 × 32 × 32 matrix, from which a submatrix of 31 × 31 × 31 was obtained. A preset threshold of 0.1 was used to convert the matrix to binary values.

The 60,000 models in the training dataset were fed to this auto-encoder with the loss function measured with cross entropy between the input models in binary values and the encode-decoded (binary) maps (see supplementary materials for details).

### 2.3 Model reconstruction from SAXS profiles

The overall workflow for model reconstruction using the auto-encoder method is as follows. First, a number of latent parameter sets (genes) are populated to get the genetic algorithm started. During each iteration, the genes are first decoded to 3D voxel objects using the auto-encoder network trained using the molecular shapes abstracted from the PISA database. The SAXS profiles are then computed using the Zernike method implemented in the SASTBX (elaborated below). Chi-scores between model profiles and target SAXS profiles are used to guide the genetic algorithm to evolve the genes until the chi-score is below a certain threshold or a pre-set number of iterations is reached. The final models are saved in density maps (CCP4 format) or bead models (PDB format).

For a voxel object, use of the 3D Zernike representation has the advantage of de-coupling the model shape and size information; thus, the SAXS profiles can be quickly evaluated if the model size is updated while the shapes are not changed. A detailed derivation was elaborated elsewhere (Liu, Morris *et al*., 2012; Liu, Hexemer *et al*., 2012), and a brief summary is provided for clarity. A 3D object *ρ*(**r**) after scaling to fit within a unit sphere can be expanded to the coefficients of 3D Zernike functions, which are a set of orthonormal polynomials with orders (n, l, m) (Canterakis, 1999; Novotni & Klein, 2003). The expansion coefficient or the so-called Zernike moment *Cnlm* is calculated with equation (1):

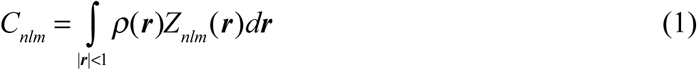

Subsequently, the 3D density distribution function *ρ*(**r**) can be approximated using {*C*_*nlm*_} up to a maximum expansion order *n*_max_.It has been shown (see (Liu, Morris *et al*., 2012)) that the SAXS intensity can be explicitly expressed as:

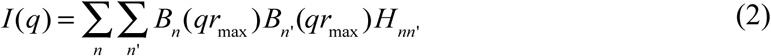

Where 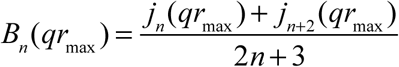 describes the contribution from corresponding Zernike polynomial to SAXS profile with the Bessel functions of the first kind *jn(x)*. In addition, the shape information is encoded in Hnn’, which is expressed using Zernike moments as follows:

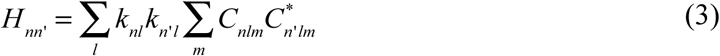

With 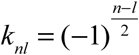. One can see that the radius of the object r_max_ is decoupled from the shape descriptors {H_nn’_}. This allows us to optimize the radius as the 201st parameter (the other 200 parameters encode the 3D shape).

The genetic algorithm was used to optimize the parameters (200 parameters for given radius information or 201 parameters if the radius is a free parameter to be optimized) (Goldberg, 1989). The target function to be minimized is the standard chi-score:

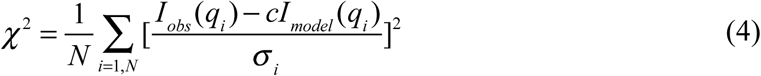

The distribution of latent parameter values obtained during the auto-encoder training procedure was used to initialize the first bunch of model parameters, so that the genes (each with 200 or 201 values) for the first generation of the genetic algorithm were populated to start the optimization. In each generation, we retain 300 genes that are generated via simulated evolution procedures. The gene evolution was implemented using three operators: selection, crossing, and mutation. The selection operator decides which genes are inherited from the previous generation by selecting the more fitted gene of two randomly selected genes from the previous generation. The crossing operator is included to exchange gene segments obtained from the previous generation. The mutation operator changes parameter values at random positions by replacing the current value to a random value with a probability distribution that follows prior knowledge of parameter distributions at that gene position.

The model comparison was measured using Pearson correlation coefficients (cc) after model alignments, which was performed using the fast rotation algorithm sastbx.superpose implemented in SASTBX (Liu, Hexemer et al., 2012). The figures are prepared using Chimera (Pettersen et al., 2004) or Pymol (Schrödinger, 2015).

## 3. Results

In this section, we first demonstrate that the auto-encoder works nicely in representing the shape information in the compressed format. Then, we show the performance of model reconstruction with or without model size information using the SAXS data as the target for optimization. The reconstructions for experimental SAXS datasets yield high-quality 3D models in general with some exceptions for challenging cases, such as loosely packed molecules or those with large cavities.

### 3.1 Quality and accuracy of auto-encoding

The voxelized objects derived from protein complex structures were encoded using 200 latent parameters in the auto-encoder neural network training procedure as described in the Methods section. From two databases for small angle scattering, the SASBDB and BIOISIS, 542 SAXS datasets with deposited 3D models were obtained (see supplementary data). First, using the 3D models from these 542 datasets, the auto-encoder performance was evaluated. Each model was converted to a 3D voxel object with binary values and then fed to the trained auto-encoder for encoding and decoding. Then, each input model was used as the reference model to assess the quality of the decoded 3D object. The real space correlation between original models and decoded models from the corresponding 200 latent parameters were computed and analysed. The auto-encoder network is very efficient and accurate in representing the majority of the 3D protein shapes, yielding a mean correlation coefficient of 0.88 with 56.1% greater than 0.90 (see Figure 1c). We also see a few failed cases, and their correlation coefficients are very low. After investigating those failed encoding cases, we found that those models have very complex shapes, such as flexible chains (for instance SASDBZ6, with cc=0.40) that are not even in a single conformation. For the majority of the testing models that have relative rigid and compact shapes, the auto-encoder works nicely in representing 3D shape information. This testing demonstrates that the 3D models are reproduced after going through the encoding-decoding procedure, and the compressed 200-d vectors are sufficient to represent 3D molecular shapes. This lays the foundation for applying the auto-encoder method to reconstruct 3D models by optimizing compressed parameters to obtain models that fit to experimental SAXS data. Furthermore, the training dataset provides the distributions of latent variables, indicating that the valid values for these variables are distributed in limited ranges (see Figure 1d for examples). This prior information will facilitate the parameter sampling during the optimization process using genetic algorithms (see Methods section).

### 3.2 Performance of reconstruction algorithm with or without model size information

The evaluation of the reconstruction algorithm was performed on the same 542 experimental datasets that were used to evaluate the performance of the auto-encoder network in the previous section. Here, we use the SAXS data as the only target to reconstruct the corresponding 3D models, and the deposited 3D structures are used as references for reconstruction model quality assessment.

The first reconstruction experiment was performed by assuming that the model size information is known. Model sizes can be derived from SAXS data using GNOM or other similar approaches (Svergun, 1992; Liu & Zwart, 2012). Instead of the maximum dimension obtained directly from the pairwise distance distribution functions, the auto-encoder method requires the radius of the model. Here, the radius of deposited models (coarse grained bead models or atomic models) from each SAXS dataset was used as an input value for model reconstruction.

The reconstruction process was monitored based on the chi-score between model SAXS profile and the experimental data. As a retrospective check, the reconstructed models are also compared with the reference model by computing their Pearson correlation coefficients after optimal alignment. Figure 2a-c present an example to demonstrate the progress of model reconstruction. The dataset is from the SASBDB (ID: SASDAH6), and the atomic structure was also deposited to allow model comparison. The average chi-scores of SAXS data comparison and the average cc of the 3D model comparison are shown for each iteration. It is clear that the chi-score was rapidly reduced within the first 10 iterations and gradually converged to a small value, indicating that the model SAXS profiles match the target SAXS data. Meanwhile, the correlation coefficients increase as the model was reconstructed. The final model SAXS profile is shown in Figure 2b compared with the experimental data (up to q=0.2 Å). The reconstructed model was superimposed onto the atomic model and shown in Figure 2c in two orthogonal orientations. The agreement between the reconstructed model (blue surface) and the atomic structure (cartoon model) illustrates the success of 3D model reconstruction for this dataset.

**Figure 2.**
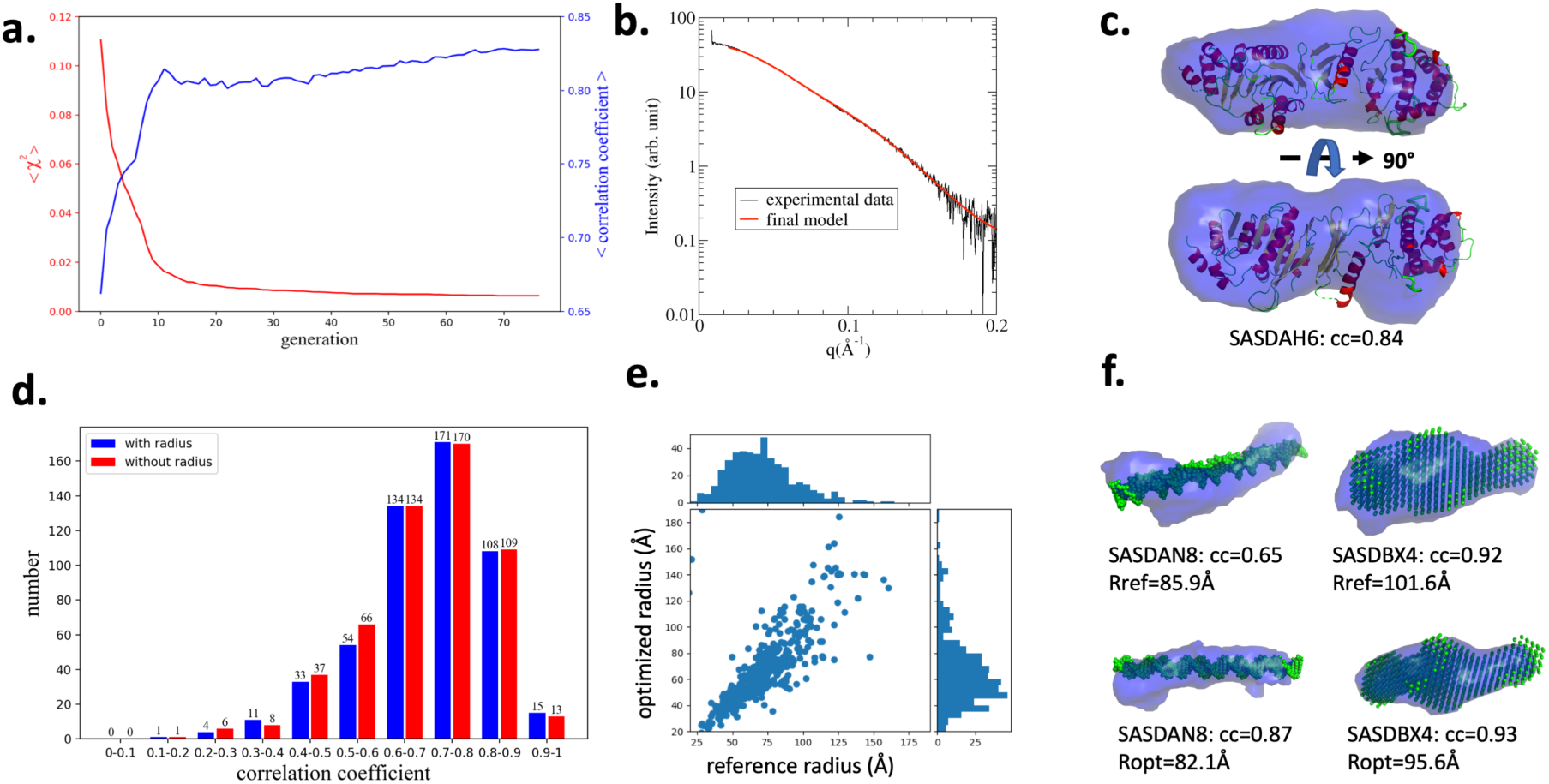
Performance of the decodeSAXS algorithm. (**a**) The progress of model reconstruction by optimizing the goodness of fit to experimental data (SASBDB ID: SASDAH6). Chi-scores for SAXS data and the Pearson correlation coefficients for reconstructed models are shown for each generation. **(b)**The SAXS comparison with experimental data for the reconstructed model in the final iteration. **(c)**The reconstructed model (blue surface) compared with the reference structure (obtained from SASBDB) at two orientations. (**d**) The algorithm performance measured using Pearson correlation coefficients between reconstructed models and the reference structures in the databases. Blue and red histograms show the statistics of correlation coefficients with or without using model size information as prior knowledge, respectively. (**e**) The comparison between optimized radii from a random value and the reference model radii. (**f**) Representative reconstructions for two examples from the SASBDB with or without radius information (more examples can be found in the supplementary materials).

Figure 2d shows the statistics of reconstructed model quality measured using Pearson correlation coefficients between the reconstructed models and the reference models in the databases. The histogram coloured in blue shows reconstruction performance using the model radius as known information. As shown in the following, the radius information is no longer required information to achieve similar accuracy levels. Among 542 testing datasets, 294 reconstructed models have correlation coefficients greater than 0.70 (see supplementary Figures for representative models at various correlation levels). At this level, the reconstructed models are consistent with the references in the overall shapes. The models with a big cavity or flexible domains are challenging for the auto-encoder to compress the shape information to a 200-dimension vector. For those models with rigid and compact structures, the auto-encoder and the SAXS-based reconstruction are very successful.

If the radius of the model to be reconstructed could not be faithfully obtained, the algorithm can optimize the size information under the same framework by simply treating the model radius as an additional element in the genes. Using the same dataset, the reconstruction algorithm was tested **without** providing accurate size information. The initial radius for each model in the first generation is a random positive number less than 300 (with the associated unit Å). The radius was taken as the 201st parameter and subject to the genetic algorithm for optimization. The optimized radii for 542 testing datasets are compared with the radii extracted from the reference models in Figure 2e, showing that the size information can be obtained by optimization. Despite radius differences in some datasets, the reconstructed model quality is comparable to the outcomes with radius information as shown in Figure 2d (red colour histogram). Two examples shown in Figure 2f demonstrate that high-quality models can be reconstructed even if the radius is not exactly the same as the reference values, indicating the robustness of the algorithm.

## 4. Discussions and Conclusion

Here, 3D model reconstruction from SAXS data is challenging given the limited information embedded in the 1D profile. Prior knowledge, especially size information, has been required for model reconstruction. Here, using the deep learning method, the 3D shape information can be compressively represented using 200-dimension vectors. More importantly, this new method does not require model size information and demonstrates its robustness in 3D model building. Currently, the 200-d vector and the SAXS profile are not directly related but indirectly related via a decoded 3D model and subsequent SAXS calculation. Preliminary results show that it is possible to encode a SAXS profile using low-dimensional vectors rather than computing from 3D models, and this feature will greatly reduce the computing time. This is the first demonstration that the deep learning method can be applied in the 3D model reconstruction from SAXS data. As more high throughput SAXS data are being collected, we anticipate increasing applications of such methods in SAXS analysis.

## Acknowledgements

Funding from National Natural Science Foundation of China (grant numbers: 11575021, U1530401, U1430237) is acknowledged.

## Notes

http://liulab.csrc.ac.cn/decodeSAXS

